# Individual Variation in Risky Decisions Is Related to Age and Gender but not to Mental Health Symptoms

**DOI:** 10.1101/2022.07.11.499611

**Authors:** Anahita Talwar, Francesca Cormack, Quentin J. M. Huys, Jonathan P. Roiser

**Author notes:** denotes corresponding author. these authors contributed equally to the manuscript (Quentin J. M. Huys, Jonathan P. Roiser).

## Abstract

Risky decisions involve choosing between options where the outcomes are uncertain. Cognitive tasks such as the CANTAB Cambridge Gamble Task (CGT) have revealed that patients with depression make more conservative decisions, but the mechanisms of choice evaluation underlying such decisions, and how they lead to the observed differences in depression, remain unknown. To test this, we used a computational modelling approach in a broad general-population sample (N = 753) who performed the CANTAB CGT and completed questionnaires assessing symptoms of mental illness, including depression. We fit five different computational models to the data, including two novel ones, and found that a novel model that uses an inverse power function in the loss domain (contrary to standard Prospect Theory accounts), and is influenced by the probabilities but not the magnitudes of different outcomes, captures the characteristics of our dataset very well. Surprisingly, model parameters were not significantly associated with any mental health questionnaire scores, including depression scales; but they were related to demographic variables, particularly age, with stronger associations than typical model-agnostic task measures. This study showcases a new methodology to analyse data from CANTAB CGT, describes a noteworthy null finding with respect to mental health symptoms, and demonstrates the added precision that a computational approach can offer.

## Introduction

Mental illnesses are often characterised by differences in decision-making, particularly in situations that involve maximising expected rewards (Cáceda et al., 2014). This has been examined extensively, with different disorders such as depression, obsessive-compulsive disorder, and psychosis showing some influence on various aspects of decision-making (Deserno et al., 2016; Halahakoon et al., 2020; Pratt et al., 2021; Sachdev & Malhi, 2005). Gambling tasks have often been used to examine such differences in decision making, requiring participants to choose between options with uncertain payoffs. These tasks are useful as they simulate the risky decisions that we often face in daily life. Early studies suggested that such tasks are sensitive to brain damage (specifically to the ventromedial prefrontal cortex, involved in fear and planning - Bechara et al., 1994). Within the realm of mental health research, prior studies using gambling tasks have suggested that individuals with depressive and anxious disorders have heightened aversion to risks and losses (Baek et al., 2017; Charpentier et al., 2017; Smoski et al., 2008), and that patients with schizophrenia make more random choices leading to lower winnings overall (Pedersen et al., 2017; Woodrow et al., 2019). However, it is difficult to integrate across such studies as they often use different paradigms, report different aspects of performance, and most patient studies have used small samples. It is also known that behaviour can be sensitive to the specific task parameter settings, such as the probabilities and gamble outcome values, even within paradigms that use largely the same framework (Peterson et al., 2021).

One particularly informative set of studies has used the Cambridge Gamble Task (CGT) from the CANTAB task battery, which requires participants to bet a proportion of their points on a simple decision. This set of studies is noteworthy because a large number of participants with a variety of diagnoses have performed the task (Ackerman et al., 2015; Deakin et al., 2004; Hutton et al., 2002; Rogers et al., 1999; Rubinsztein et al., 2001). The CGT was originally designed to remove some of the learning confounds present in previously popular gambling tasks, such as the Iowa gambling task (Bechara et al., 1994), and presents participants with explicit information about the values and probabilities of gambles (Rogers et al., 1999). One of the most consistent findings with the CGT in mental health research is that, relative to controls, depressed individuals choose to bet fewer points overall, particularly when the probability of winning is high (Mannie et al., 2015; Murphy et al., 2001; Rawal et al., 2013). This has usually been interpreted as reflecting a conservative, or risk-averse, decision-making strategy. Studies observing this pattern have included diverse groups including adult patients with depression (Murphy et al., 2001), young people with a family history of depression but who had not been diagnosed themselves (Mannie et al., 2015), and adolescents with depression (Rawal et al., 2013). However, this pattern was not observed in a smaller study including patients with bipolar depression (Rubinsztein et al., 2006), and the opposite pattern was found in a study that included adolescents with recent first episode depression, with patients betting more overall than controls (Kyte et al., 2005). In addition to the small sample sizes in these studies, and the different groups examined, another possible explanation for these inconsistent results is that the dependent variables typically examined are multifactorial. For instance, the ‘overall proportion bet’ measure depends on the proportion of points that a participant chose to bet on trials over the entire task, and disregards the specific aspects of different trials, such as the probability of winning or the stake. This challenges the interpretation of results as the key underlying mechanistic processes cannot be examined.

Computational models of cognitive neuroscience tasks are promising due to their ability to account for trial-by-trial behaviours, dissect traditional measures into more precise components, and make concrete predictions about participants’ choices at an individual level. The most well-known example of a computational model applied to decision-making is Kahneman and Tversky’s Prospect Theory, which summarises performance in terms of parameters such as risk aversion (the degree to which participants avoid uncertainty) and loss aversion (the degree to which losses loom larger than gains) (Kahneman & Tversky, 1979). These types of models have been applied in mental health research where it has been shown that anxious patients are more risk averse, but not more loss averse, than healthy controls (Charpentier et al., 2017). Computational models based on Prospect Theory have also been applied to the CANTAB CGT. Romeu and colleagues modified traditional models to better fit CGT data from controls and groups of patients with various substance use disorders (Romeu et al., 2020). They found that whilst standard task measures such as “quality of decision making” (reflecting the tendency to make high-probability choices) and “risk adjustment” (the calibration between betting behaviour and probability), showed little difference between patients and controls, model parameters such as risk sensitivity and delay aversion did vary between groups, highlighting the added precision that the modelling approach can provide. However, in this study, the authors did not show some important aspects of model checking, such as parameter recovery and correlations between individual-level summary measures (such as ‘overall proportion bet’) from model-simulated and real data. In particular, the latter is more specific than posterior predictive checks on the average of group behaviour and is important to highlight potential areas of model weakness (Wilson & Collins, 2019). Additionally, in this prior work the authors focused largely on capturing the impulsivity aspect of task performance due to its relevance in substance use disorders; however, mood and anxiety disorders are hypothesised to be more closely associated with changes in appetite for risk.

Here, we expand on this prior work by combining several approaches. First, we examine CGT behaviour in a large sample with a number of self-reported measures of psychopathology. This allows us to jointly assess the relationship between computationally defined decision-making processes, psychopathology and demographic variables. Second, we build on a previous computational approach (Romeu et al., 2020) to develop a model which is fully validated. Third, we collect data online which enables fast scaling and replication in novel cohorts. The validated model can then be applied to existing datasets to understand the relationship between any group or psychopathology measures and well-defined computational processes.

## Materials and Methods

### Participants

Our dataset was collected online via Prolific Academic. Participants were recruited if they confirmed that they: a) were over 18 years of age; b) were fluent in English; c) had not experienced a significant head injury (resulting in loss of consciousness); d) had not been diagnosed with an untreated mental health condition (by medication or psychological intervention) that had a significant impact on their daily life; e) had never been diagnosed with mild cognitive impairment or dementia. All participants were paid at a rate of £7.50 per hour. Participants were anonymous, and they provided informed consent online before participating in the experiment. The dataset includes 762 participants who completed the CGT task and also several self-report mental health questionnaires. This sample size provides 95% power to detect associations of r = 0.13 at alpha = 0.05 (two-tailed).

### CANTAB Cambridge Gamble Task

Participants completed the CANTAB CGT, as described previously (Rogers et al., 1999). The design of the task is presented in Figure 1A. Participants start with 100 points and are presented with ten boxes at the top of the screen on each trial - some of which are red, and the rest blue. They are told that one box has a token in it and that they must guess the colour of the box containing the token by selecting their choice at the bottom of the screen. They then have to bet a proportion of their points that their guess is correct. The possible bets (0.05, 0.25, 0.50, 0.75 or 0.95 of their current points) are presented in a circle in the centre of the screen for two seconds each. Participants click on the circle when they see the amount they want to bet. The result of their choice is shown, and if they are correct, the points are added to their total. If not, the points are deducted. A new trial begins with different numbers of red and blue boxes.

**Figure 1.**
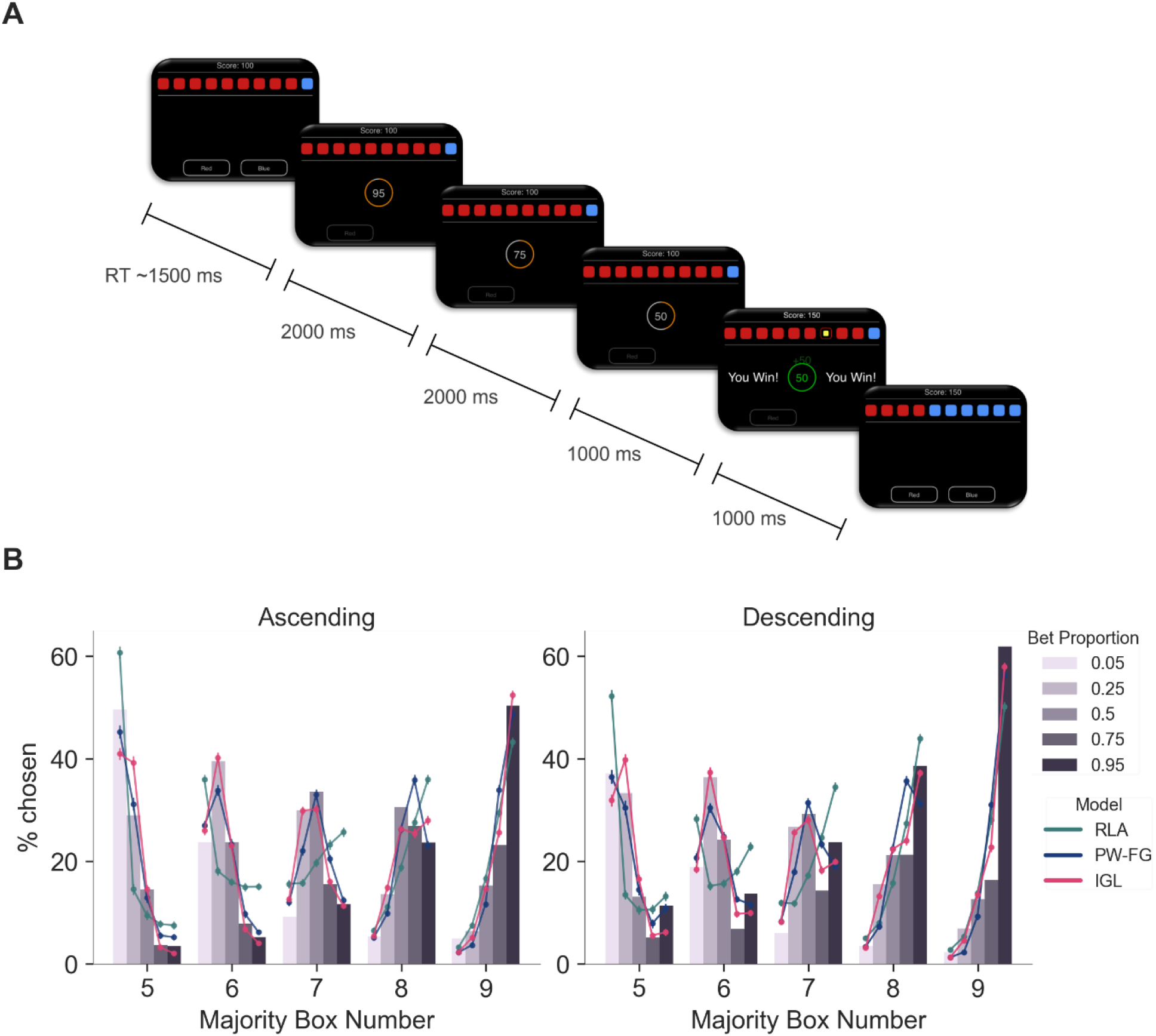
Task and Betting Behaviour. **A**. Schematic of a trial from the descending condition of the CANTAB CGT. Red and blue boxes are displayed at the top of the screen. After participants select which colour box they think the token is behind, bet options (0.95, 0.75, 0.50, 0.25, 0.05 of their current points – here 100) are presented sequentially in the centre of the screen. Participants click on the number they want to bet before the token is revealed. The points bet are added or subtracted from the total depending on whether the colour choice was correct or incorrect, and a new trial begins with a new box ratio and the new points total. **B**. Summary of real and model-simulated betting behaviour. Plots show the percentage of trials with each majority box number in which participants chose each bet proportion. Real data are shown with bars, while model-simulated data is shown by the coloured lines. Points show the mean from 100 simulations, and error bars depict the standard deviation. Data are shown separately for the ascending (left) and descending (right) task conditions, and only trials on which participants chose the majority box colour (and all 5:5 trials) are included. RLA – Risk and Loss Aversion; PW-FG – Projected Wealth Fixed Gains, IGL – Inverse Gains and Losses.

Participants completed 8 practise trials in which they first completed the colour choice part of the task on its own, before the bet component was added; these trials were not included in task analysis. For the first 18 assessed trials, the stakes were shown in descending order, and for the subsequent 18 trials, in ascending order. Participants were excluded from analysis if they did not attempt all four blocks of the task (n = 2), selected either the earliest or the latest bets on all trials (n = 4), or selected the non-majority box colour on more than 50% of trials (n = 3) leaving 753 participants for modelling analysis.

### Self-Report Questionnaires

Participants provided their age, gender and level of education^1^. They also completed questionnaires assessing depressive symptoms (Self-Rating Depression Scale, SRDS (Zung, 1965)), anxious symptoms (State Trait Anxiety Inventory, STAI (Spielberger, 1983)), impulsivity (Barratt Impulsiveness Scale, BIS-11 (Patton et al., 1995)), and anhedonia (Temporal Experience of Pleasure Scale, TEPS (Gard et al., 2006)).

### Descriptive Measures of CGT Performance

CANTAB CGT data are typically analysed using a number of descriptive (or model-agnostic) measures (Deakin et al., 2004; Rogers et al., 1999), which we detail here as they are used to assess model performance throughout the paper:

1. ***Quality of Decision Making (QDM)*** The proportion of trials on which the participant chose the majority box colour, calculated over all trials on which the number of boxes in each colour differed.

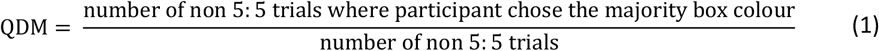
2. ***Overall Proportion Bet (OPB)*** The mean proportion of current points gambled by the subject on all gamble trials.

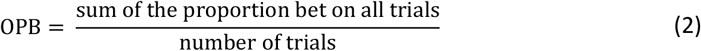
3. ***Risk Adjustment (RA)*** A measure of a participant’s sensitivity to probability when betting. Higher values suggest that participants increase their bets with increasingly favourable odds (box ratio), while lower values suggest participants bet consistently irrespective of box ratio. This includes trials on which participant chose the majority box colour and all 5:5 trials.

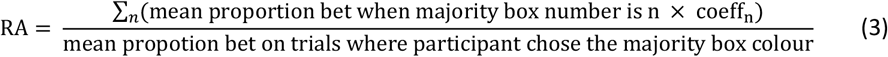

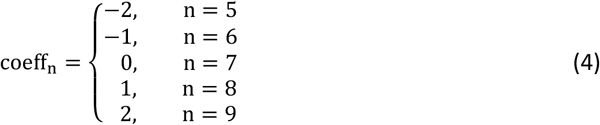
4. ***Delay Aversion (DA)*** The difference between the mean proportion bet in the ascending and descending conditions. This includes trials on which participant chose the majority box colour and the 5:5 trials.

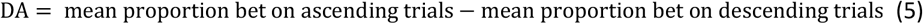

### Computational Models

While the above descriptive measures represent intuitive summary statistics of task performance, they do not provide insight into *why* participants are making the choices they are. Computational models, on the other hand, use a generative approach to specify how participants interpret the stimuli presented to them and use information to make decisions, and therefore capture the underlying cognitive mechanisms involved in completing the task. Thus, we developed computational models to capture trial-by-trial decision making and risk-taking processes to directly test which processes can account for individual differences in gambling task behaviour.

Each trial of the task requires two decisions - first to choose the box colour, and second to choose what proportion of points to gamble. The likelihood for each trial combines the model’s predictions of these two choices in the following way: *p(chosen colour)* × *p(bet*|*chosen colour)*, such that the bet choice is dependent on the prior colour choice.

All models make the same assumptions about the probability of choosing colours. Let *c*_*t*_ be the choice of colour on trial t, then:

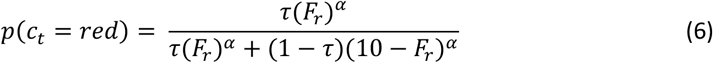

Where *F*_*r*_ is the number of red boxes on the screen on the current trial (range 0 - 10), τ is a red-bias parameter such that higher values mean the participant is more biased to selecting red, and *α* is a sensitivity parameter that indicates how sensitive participants are to the box ratio on each trial, equivalent to the slope of a logistic function. Higher values of *α* indicate that the participant chooses the majority box colour in a more deterministic manner.

The models then compute the probability of bets *p*(*b*_*t*_|*c*_*t*_). The first step is the evaluation of the utility *U* of each bet option. This is where the different models differ, and we will turn to the different formulations of this below. Next, the utility of each potential outcome (a win or a loss of a certain magnitude, or the future wealth) is weighted by the probability of that outcome based on the choice of colour *c*_*t*_. Finally, a linear delay factor is included which penalizes options that are presented later. This results in an overall value *V*(*b*_*t*_|*c*_*t*_) for each bet *b*_*t*_, conditioned on the first-stage colour choice *c*_*t*_ as follows:

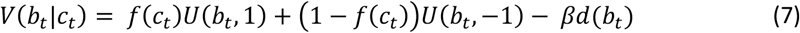

where the function *d*(*b*_*t*_) takes on values {0, 0.25, 0.5, 0.75, 1} for increasing delays *d*() of the bets, and the function *f*(*c*_*t*_) indicates the fraction (number of boxes/10) of the chosen colour on that particular trial. The form and meaning of *U* depends on the particular model as explained further below. Finally, the values *V* determine the probabilities of choosing each bet through a softmax function:

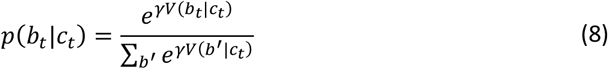

and we consider the joint probability of colour and bet choices: *ℒ*(*θ*) = ∏_*t*_ *p* (*c*_*t*_|*θ*)*p*(*b*_*t*_|*c*_*t*_, *θ*) for inference of parameters *θ*.

#### 1. Risk and Loss Aversion (RLA)

The first model is based on the influential framework of Kahneman and Tversky (Kahneman & Tversky, 1979). Prospect Theory assumes that participants subjectively value the potential wins and losses on each trial. In the CGT, participants bet a proportion of their points which are either added or subtracted from their total, so the potential wins and losses are a product *b*_*t*_*w*_*t*_ of the bet *b*_*t*_ chosen on trial *t*, and the wealth *w*_*t*_ on that trial. The bet takes on a fixed set of proportions (*b*_*t*_ ∈ .05, .25, .5, .75, .95). In the RLA model, the subjective utility of the wins and losses is defined as follows:

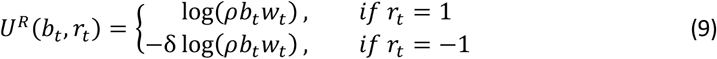

where *r*_*t*_ = 1 indicates that the colour choice was successful, *r*_*t*_ = −1 that it was not. The parameter *ρ* scales the wins and losses, and functions as a risk-aversion parameter because it is placed inside the nonlinear logarithm function, and hence alters the effective shape of the utility curve. δ is a loss-aversion parameter, such that large δ means that losses are more aversive than the equivalent gain is rewarding. Note that we have used a log function (as opposed to the conventionally used power function) to aid numerical stability.

The parameters in this model were *θ* = {*α, c, ρ, δ, β, γ*} where *α* is the colour choice determinism, *c* the colour choice bias, *ρ* the risk aversion, *δ* the loss aversion, *β* the delay aversion and *γ* the bet choice determinism.

#### 2. Projected Wealth (PW)

The RLA model as formulated above considers the potential wins and losses that would result on each trial. We next consider a related model (Bernoulli, 1738; von Neumann & Morgenstern, 1944) which assumes that the attractiveness of different bet options depends on the total projected wealth each option would result in, *w*_*t*_ + *b*_*t*_*w*_*t*_ *r*_*t*_. The utility *U*^*w*^ of each bet in the wealth model is defined as:

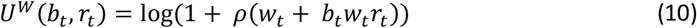

Note that this model does not include a loss aversion parameter. The parameters in this model were *θ* = {*α, c, ρ, β, γ*} where *α* is the colour choice determinism, *c* the colour choice bias, *ρ* the risk aversion, *β* the delay aversion and *γ* the bet choice determinism.

#### 3. Projected Wealth Fixed Gains (PW-FG)

We next investigated the winning model from Romeu et al., which is similar to the PW model with the exception that the risk aversion parameter in the domain of gains is fixed to 1, and the parameter is only estimated in the loss domain. Again, we assume that participants subjectively value the potential total wealth, *w*_*t*_ + *b*_*t*_*w*_*t*_ *r*_*t*_.

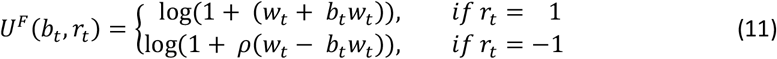

The parameters in this model were *θ* = {*α, c, ρ, β, γ*} where *α* is the colour choice determinism, *c* the colour choice bias, *ρ* the risk aversion, *β* the delay aversion and *γ* the bet choice determinism.

#### 4. Linear Loss Aversion (LLA)

We developed further models in order to improve the performance compared to the previously published PW-FG model described above. Our first novel model assumes that participants subjectively value the bet proportions *b*_*t*_, independently from their current wealth (note that unlike in the above models, *w*_*t*_ does not enter the specification). Gains can be distorted through a power function (representing risk aversion), while losses are scaled linearly (representing loss aversion).

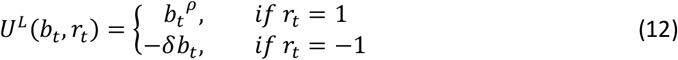

The parameters in this model were *θ* = {*α, c, ρ, δ, β, γ*} where *α* is the colour choice determinism, *c* the colour choice bias, *ρ* the risk aversion, *δ* the loss aversion, *β* the delay aversion and *γ* the bet choice determinism.

#### 5. Inverse Gains and Losses (IGL)

Our second novel model also assumes that participants subjectively value the bet proportions *b*_*t*_, again independently from their current wealth.

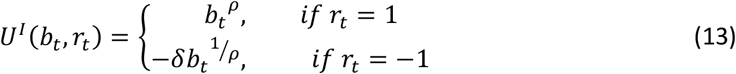

In this model, the distortions of loss and gains are linked by an inverse power function, such that if one is convex, the other is too, and if one is concave, the other is too. This breaks the assumption of Prospect Theory of risk aversion in the gains domain, combined with risk seeking in the losses domain. Losses are again additionally linearly scaled by a loss aversion parameter *δ*.

The parameters in this model were *θ* = {*α, c, ρ, δ, β, γ*} where *α* is the colour choice determinism, *c* the colour choice bias, *ρ* the risk aversion, *δ* the loss aversion, *β* the delay aversion and *γ* the bet choice determinism.

### Parameter Estimation

As in Talwar et al., 2021, we used a hierarchical Bayesian parameter estimation approach, described previously (Huys et al., 2011), which finds the maximum *a posteriori* parameter estimates for each participant, given the model and the data, and sets the parameters of the prior distribution to the maximum likelihood estimates given all participants’ data. The purpose of using hierarchical estimation here is primarily that priors over the parameters act to regularise the estimates such that unrealistic, extreme values are avoided. We used an expectation-maximisation approach which repeatedly iterates over two steps until convergence is reached. Briefly, in the E-step the model finds the best-fitting individual level parameter estimates for each participant given their data and the current parameters of the prior distribution, and in the M-step, the maximum likelihood group level prior parameters are updated to reflect the current individual parameter estimates.

All parameters were transformed to ensure that 0 ≤ *c* ≤ 1, and *α, ρ, δ, γ* ≥ 0. The mean and standard deviation for the best-fitting parameter values of each model are given in Table 1. Recoverability of parameters was calculated by simulating 300 datasets with parameters drawn randomly from the estimated prior distribution. The best-fitting parameters for these data sets were found as above, and parameter recoverability is indicated by the correlation between the simulated and recovered parameters (Table 1). Untransformed parameter values were used for statistical inference as these are estimated to be distributed with a multivariate Gaussian distribution.

**Table 1.** Free parameters of each model, along with summary statistics of their best-fitting values and recoverability.

| Model | Parameter | Range | Mean ± SD, Median | Recovery (r) |
| --- | --- | --- | --- | --- |
| 1. Risk and Loss Aversion | Colour choice determinism | 0 - ∞ | 8.30 ± 4.76, 8.48 | 0.71 |
| | Colour choice bias | 0 - 1 | 0.50 ± 0.07, 0.50 | 0.50 |
| | Risk aversion | 0 - ∞ | 0.86 ± 0.01, 0.86 | 0.00 |
| | Loss aversion | 0 - ∞ | 1.90 ± 1.13, 1.67 | 0.87 |
| | Delay Aversion | $-\infty$ - ∞ | 0.22 ± 0.46, 0.15 | 0.85 |
| | Bet choice determinism | 0 - ∞ | 3.19 ± 3.00, 2.50 | 0.89 |
| 2. Projected Wealth | Colour choice determinism | 0 - ∞ | 8.38 ± 4.81, 8.58 | 0.62 |
| | Colour choice bias | 0 - 1 | 0.51 ± 0.07, 0.50 | 0.57 |
| | Risk aversion | 0 - ∞ | 0.98 ± 8.06, 0.02 | 0.00 |
| | Delay Aversion | $-\infty$ - ∞ | 0.02 ± 0.04, 0.02 | 0.77 |
| | Bet choice determinism | 0 - ∞ | 33.27 ± 30.87, 24.15 | 0.86 |
| 3. Projected Wealth Fixed Gains | Colour choice determinism | 0 - ∞ | 8.48 ± 5.11, 8.38 | 0.69 |
| | Colour choice bias | 0 - 1 | 0.51 ± 0.07, 0.51 | 0.61 |
| | Risk aversion | 0 - ∞ | 0.13 ± 0.21, 0.03 | 0.51 |
| | Delay Aversion | $-\infty$ - ∞ | 0.05 ± 0.13, 0.04 | 0.93 |
| | Bet choice determinism | 0 - ∞ | 18.47 ± 10.60, 16.23 | 0.89 |
| 4. Linear Loss Aversion | Colour choice determinism | 0 - ∞ | 8.16 ± 4.53, 8.52 | 0.58 |
| | Colour choice bias | 0 - 1 | 0.50 ± 0.07, 0.50 | 0.53 |
| | Risk aversion | 0 - ∞ | 9.88 ± 20.06, 0.44 | 0.16 |
| | Loss aversion | 0 - ∞ | 1.29 ± 1.14, 0.99 | 0.43 |
| | Delay Aversion | $-\infty$ - ∞ | 0.08 ± 0.09, 0.04 | 0.71 |
| | Bet choice determinism | 0 - ∞ | 38.98 ± 35.68, 29.16 | 0.43 |
| 5. Inverse Gains and Losses | Colour choice determinism | 0 - ∞ | 8.38 ± 5.01, 8.55 | 0.62 |
| | Colour choice bias | 0 - 1 | 0.50 ± 0.07, 0.50 | 0.53 |
| | Risk aversion | 0 - ∞ | 0.68 ± 0.35, 0.62 | 0.85 |
| | Loss aversion | 0 - ∞ | 2.65 ± 2.36, 1.99 | 0.95 |
| | Delay Aversion | $-\infty$ - ∞ | 0.07 ± 0.16, 0.05 | 0.91 |
| | Bet choice determinism | 0 - ∞ | 18.47 ± 12.62, 15.65 | 0.80 |

### Model Comparison

As in Talwar et al., 2021, *a priori* we did not assume that any of the models should be more likely. Therefore, they can be compared by examining the approximate model log likelihood with the integrated Bayesian Information Criterion (Huys et al., 2011). This procedure gives us the model that fits the data most parsimoniously, whilst penalising for unnecessary added model complexity (additional parameters).

We carried out posterior predictive checks (i.e. qualitative model comparisons) in which we assessed model performance by comparing model-simulated data, using participants’ estimated parameters, to real data. This involved comparing overall group level performance patterns, and correlating individual task summary statistics. For the latter, ten simulated datasets were run for each participant and correlations with real data were calculated for each iteration. The average correlation is reported as a metric of model fit.

### Sensitivity Analysis

We carried out a sensitivity analysis by removing participants whose bet choice data was not fit better by our winning model compared to chance performance. For this, we counted the number of times the model assigned the greatest probability to the bet that the participant actually chose and used a binomial test to compare this to the number of times we would expect it to happen by chance (where the probability of choosing each bet proportion was 1/5). We kept all participants where the outcome of this test was significant (suggesting that the winning model provided a significantly better fit to their data than chance) leaving a sample size of 727 (N = 26/753 (3.5%) excluded) for this analysis.

## Results

Our final dataset consisted of 753 participants with an age range of 18 – 76 years (mean = 41.5, sd = 13.5), and of which 358 (50%) were women. In Figure 1B, the bars show the betting patterns in the raw data - people tend to pick smaller bets when the box ratio is more evenly matched (lower majority box number) but bet more points when there is greater discrepancy in the box colours (higher majority box number). Furthermore, there is a bias towards betting higher values in the descending condition compared to the ascending condition (descend vs ascend: t(752) = 9.77, p = 2.63×10^−21^), suggestive of delay averse (impatient) behaviour.

Inspired by the modelling framework in Romeu et al., 2020, all our models had the same colour choice (Eq 6), probability weighting (Eq 7), delay aversion (Eq 7), and bet choice determinism (Eq 8) functions, while the key distinction between models came from the valuation functions (Eq 9 - 13).

### Previous Models of Risky Decision Making Perform Poorly

We first assessed the performance of three previously published models for risky decision making. The first valuation function we used was inspired by the influential Prospect Theory (Kahneman & Tversky, 1979). The Risk and Loss aversion model (RLA) proposes that participants subjectively evaluate the amount that could be won or lost on each trial with marginally decreasing utility (i.e. risk aversion), that people are loss averse (losses are more aversive than gains are attractive), and that people are risk seeking in the domain of losses, but risk averse in the domain of gains. In the behavioural economics literature, these models are typically fit to group behaviour and the average parameter estimates obtained. Here, we are more interested in fitting individual-level rather than group-level parameters to explore how differences in e.g., loss aversion and risk aversion lead to differences in task behaviour. When examining the predictions made by the RLA model, we found that it predicts that participants were most likely to choose either the largest or the smallest bet available on all trials. The green line in Figure 1B shows how the model predicts that the 0.05 or 0.95 bets are chosen the most often regardless of the box ratio, in contrast to the human data which displays a clear preference for intermediate bets when the majority box number is between 6 - 8. Therefore, the RLA model was not able to capture the key betting patterns observed in the human data (Figure 1B, S1A).

The second established model we tested was inspired by Expected Utility Theory (Bernoulli, 1738; von Neumann & Morgenstern, 1944). The model assumes that participants subjectively evaluate their projected wealth, rather than losses and gains, and therefore does not include a loss aversion parameter. Unlike the RLA model, this Projected Wealth model (PW) model was able to generate participants’ choosing intermediate bet proportions (0.25, 0.50, 0.75). However, the model was not able to capture the behaviour of the most low-betting participants (Figure S1B).

The Projected Wealth Fixed Gains model (PW-FG; Romeu et al., 2020) is a previously published model that was specifically developed to describe behaviour on CANTAB CGT. It is similar to the PW model except for the adaptation that the risk aversion parameter was fixed to 1 for potential increases in cumulative points while a free parameter was estimated for the case of potential decreases to cumulative points. However, the PW-FG model exhibited the same weakness as the PW model, failing to account for the behaviour of participants who chose lower bets on average, even at high box ratios, as shown most clearly in Figure 1B. In particular it overestimates the percentage of participants that chose to bet 75% of their points when the majority box number was 7, 8 or 9. This weakness of the PW-FG model is further emphasised by the scatterplot of the overall proportion bet measure in real vs model simulated data, where the failure to capture the behaviour of low-betting participants is clear (Figure S1C). Capturing this conservative betting behaviour is of particular importance as prior studies indicate that placing low bets on favourable gambles is related to anxious and depressive symptoms (Charpentier et al., 2017; Murphy et al., 2001; Rawal et al., 2013).

### Participants are Risk Averse for both Gains and Losses, and Indifferent to Current Wealth

In order to overcome the limitations of the three models described above, we developed two novel models in which participants are assumed to subjectively evaluate the bet proportions themselves (0.05, 0.25, 0.50, 0.75, 0.95) thus rendering their current number of points irrelevant for decision making. The Linear Loss Aversion model (LLA) uses a power function for gains, and a linear function was introduced for losses to prevent the convexity that led to the RLA model being unable to predict participants’ choosing intermediate bets. Further, a loss aversion parameter was reincorporated to overcome the limitation of the PW and PW-FG models, and better describe the behaviour of consistently low-betting participants. This model performed much better than previous ones as shown by the promising correlation between real and model-simulated data, particularly for risk adjustment scores (Figure S1D). We then developed a final model to improve the model’s predictions of participant-specific risk adjustment further, as this is a key feature of task performance. This model, Inverse Gains and Losses (IGL), also uses a power function for gains with the subjective valuation of losses now also determined by a power function and parameterised with the inverse risk aversion parameter (1/ρ). This has the effect that predicted behaviour is consistent between the domains of gains and losses, being either risk-seeking or risk-averse in both, in contradiction to the RLA model. This model was able to fully recapitulate the patterns of individual variability observed (Figure 1B).

Despite the utility of posterior predictive checks (Figure 1B, S1) during our iterative model development procedure, a more formal model comparison approach is required. Figure 2A shows the average likelihood per trial, an indication of how well the model predicts participant choices on average, as well as the integrated Bayesian Information Criterion, which considers both the model’s predictions and simultaneously penalises for added model complexity (a less negative score indicates a better model). These graphs both show that each successive model improves overall model performance, with the Inverse Gains and Losses model performing best on both measures. Figure 2B demonstrates how this winning model generates betting choices based on the box ratio presented to the participant on a particular trial, using the group median of estimated parameter values. Of note is the concavity of the valuation of the bet proportions in the far-left panel, which suggests that participants are indeed largely risk averse in the domain of both gains *and* losses, and the quite minimal effect of delay aversion which is evident by comparing the middle two panels. Finally, Figure 3 shows that the model is able to capture both the group-level behaviour (3A), and the individual-level descriptive measures (3B) very well, as the correlations with the descriptive measures are all 0.89 and above.

**Figure 2.**
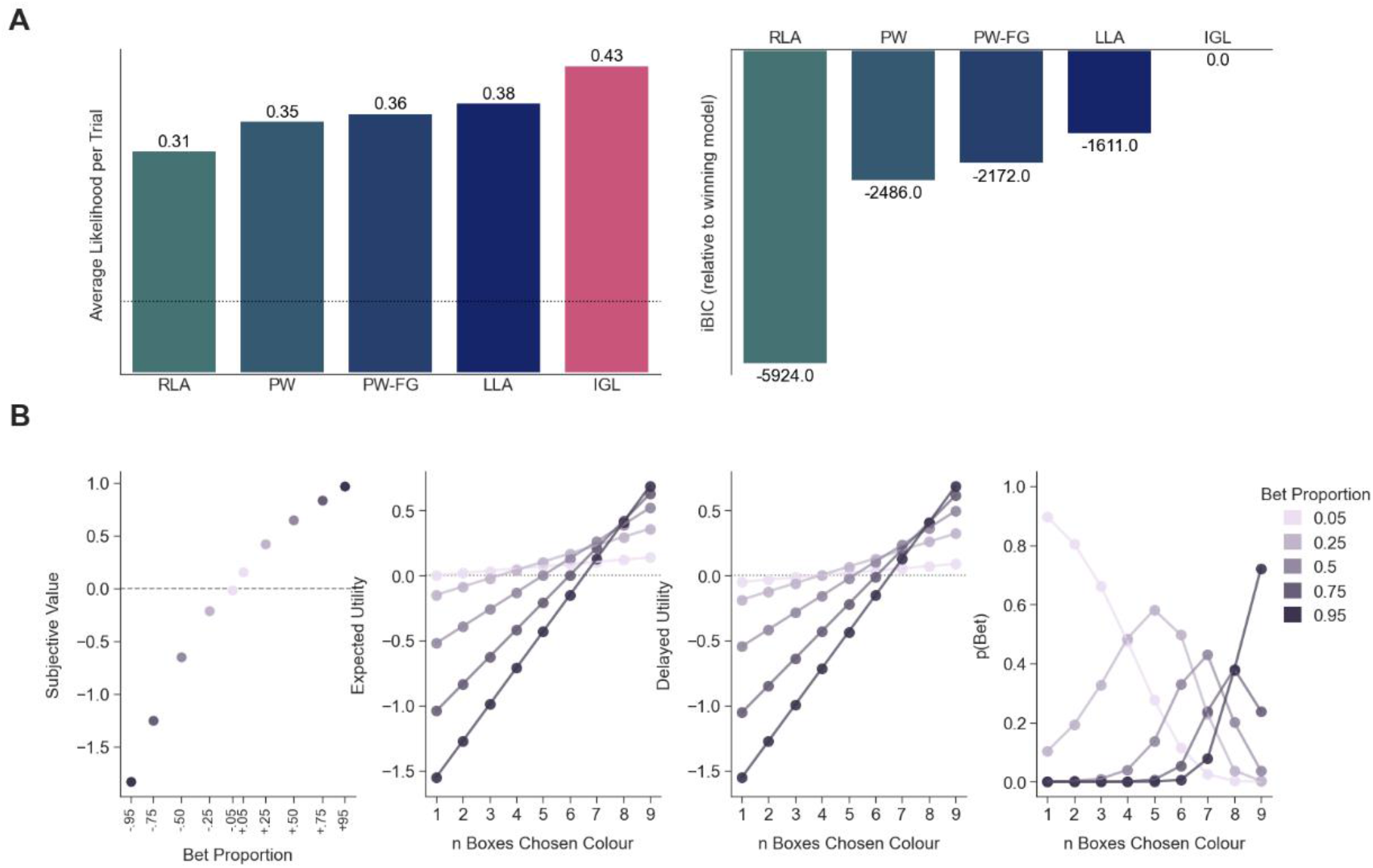
Model Comparison and Model Simulations. **A**. Model Comparison. Model performance assessed by average likelihood per trial (left) and integrated Bayesian Information Criterion (right). The dotted line indicates the average likelihood per trial for a model that makes choices at random – 0.1. **B**. Model Simulations. Internal model values simulated using the best fitting Inverse Gains and Losses (IGL) model. *Far Left:* Subjective valuation of potential wins and losses for each bet proportion. *Centre Right:* Win/loss values are weighted by their probabilities (n boxes of the chosen/unchosen colour) to give the expected utility for each bet proportion. *Centre Left:* Expected utilities are adjusted by the order in which they are displayed. This example is from the descending condition such that lower bet proportions are shown later, and thus penalised more. *Far Right:* The delayed utilities are passed through a softmax equation to give the probability of a participant choosing each bet. Importantly the IGL model predicts that participants are likely to make intermediate-level bets when the box ratio is between 6-8, consistent with real behaviour (cf Figure 1B). The medians of estimated parameters were used for simulations: risk aversion – 0.62, loss aversion – 1.99, delay aversion – 0.05, bet choice determinism – 15.65. The dotted lines at 0 aid visualisation. RLA – Risk and Loss Aversion; PW – Projected Wealth, PW-FG – Projected Wealth Fixed Gains, LLA – Linear Loss Aversion, IGL – Inverse Gains and Losses.

**Figure 3.**
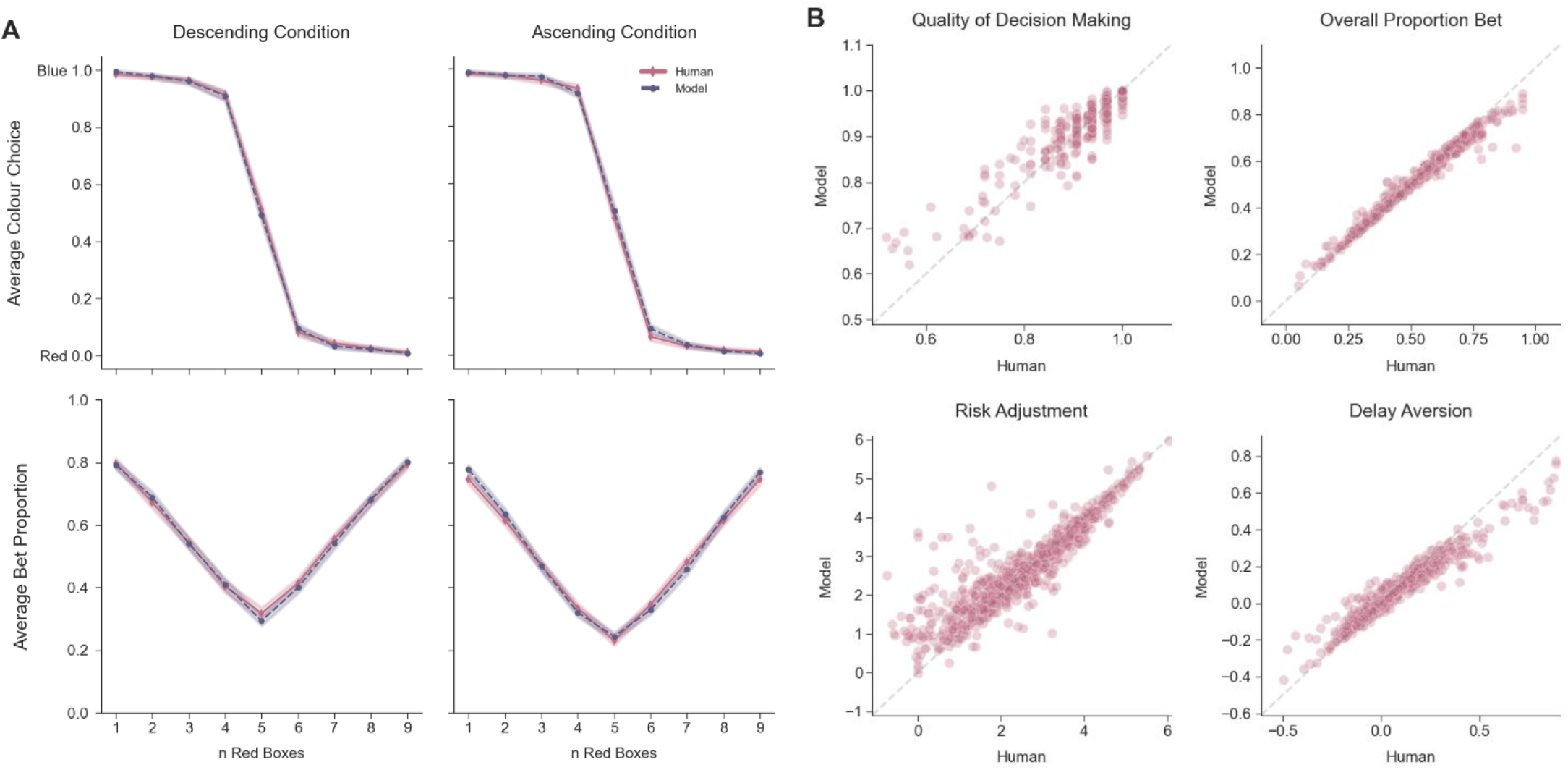
Qualitative Fits for Inverse Gains and Losses Model. **A**. Average Group Fits. Average participant and model-simulated patterns of colour choice (top) and proportion bet (bottom) for each box ratio. Points indicate mean and bands represent 95% confidence intervals. **B**. Individual Differences of Descriptive Measures. Scatter plot of human vs model-simulated scores for Quality of Decision Making, r = 0.94 (top left), Overall Proportion Bet, r = 0.99 (top right), Risk Adjustment, r = 0.89 (bottom left), and Delay Aversion, r = 0.96 (bottom right). The dotted line indicates y = x (perfect prediction). The correlation between human and model data was calculated for 10 different simulations, and the average of these is reported here.

### Risk Aversion Increases with Age, and Women are More Risk Averse Than Men

To assess which model parameters were related to age, gender and level of education, we calculated Pearson’s correlations and t-tests. Age was negatively associated with risk aversion and delay aversion, but positively associated with loss aversion (risk aversion: r(751) = -0.24, p = 1.49×10^−11^; delay aversion: r(751) = -0.09, p = 0.011; loss aversion: r(751) = 0.10, p = 0.0049). Women were less deterministic in both the colour choice and betting choice part of the task and were also more risk-averse and loss-averse than men (colour choice determinism: t(751) = 2.82, p = 0.0049, Cohen’s d = 0.21; bet choice determinism: t(751) = 2.71, p = 0.0069, Cohen’s d = 0.20; risk aversion: t(751) = 5.03, p = 6.23×10^−07^, Cohen’s d = 0.37; loss aversion: t(751) = 2.45, p = 0.015, Cohen’s d = -0.18). Finally, higher education level was positively correlated with colour choice determinism (r(751) = 0.08, p = 0.027). After applying a Bonferroni correction to account for the 18 comparisons performed (α = 0.0027), the relationships between risk aversion and age/gender remained significant (Figure 4).

**Figure 4.**
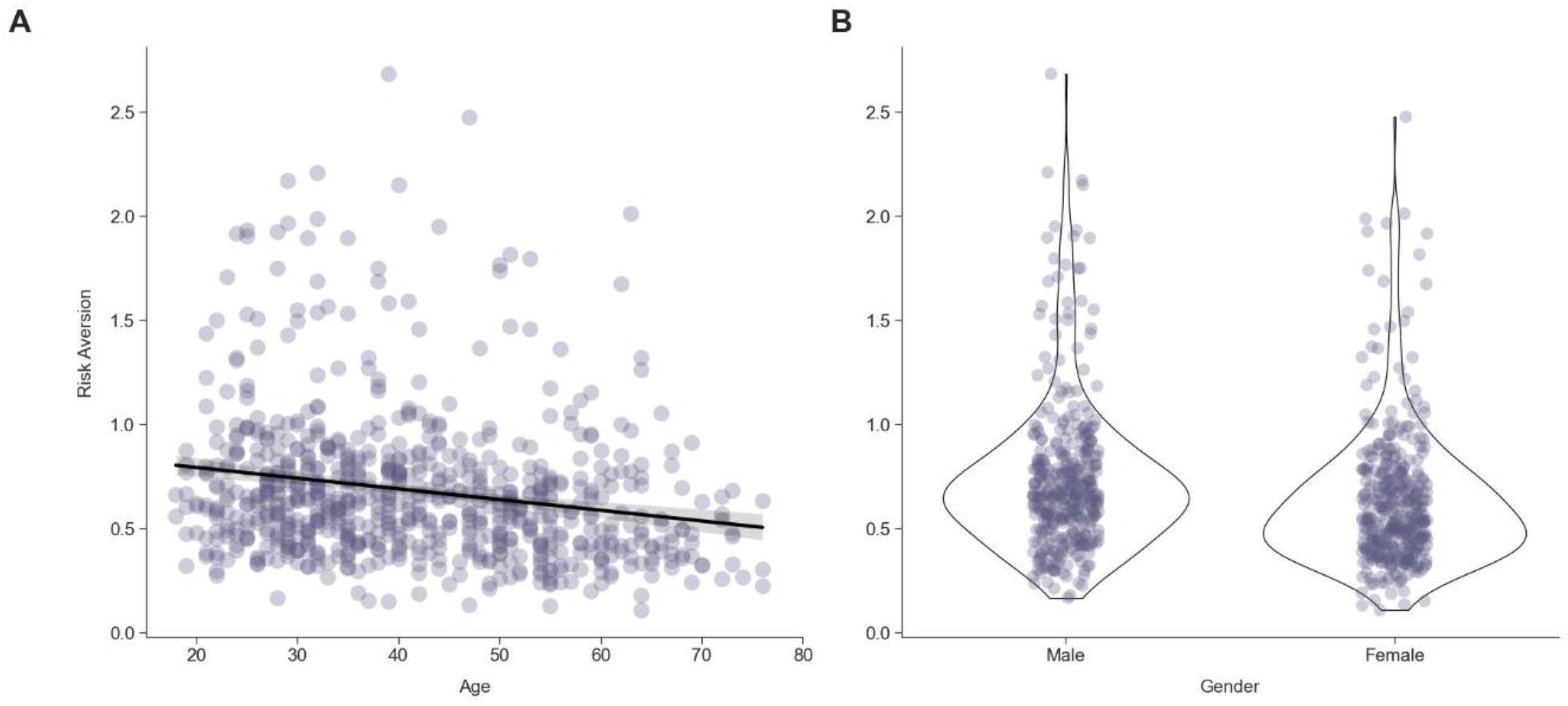
Associations Between Parameters and Demographic Variables. **A**. Older people are more risk averse. Lower values of the risk aversion parameter represent higher risk aversion, r(751) = -0.24. **B**. Women are more risk averse. Lower values of the risk aversion parameter represent higher risk aversion, t(751) = 5.03. Untransformed parameter values are used for statistical inference while transformed values (*ρ* > 0) are used for visualisation.

Age and gender were also associated with the model-agnostic measures: age was negatively correlated with overall proportion bet, risk adjustment and delay aversion (overall proportion bet: r(751) = -0.14, p = 1.07×10^−4^; risk adjustment: r(750) = -0.11, p = 3.80×10^−3^; delay aversion: r(749) = -0.087, p = 0.017) whilst women bet less, and adjusted their bets less compared to men (overall proportion bet: t(751) = 4.13, p = 4.09×10^−5^, Cohen’s d = 0.30; risk adjustment: t(750) = 3.51, p = 4.72×10^−4^, Cohen’s d = 0.26)^2^. After applying a Bonferroni correction to account for the 12 comparisons performed (α = 0.0042), the relationships between overall proportion bet and risk adjustment with both age and gender remained significant. Crucially, the size of the relationship between age and the computational risk aversion parameter was significantly greater than that with the model-agnostic overall proportion bet measure (r = -0.24 vs r = -0.14: Steiger’s Z = 2.48, p = 0.0066), demonstrating the extra sensitivity conferred by the computational approach. The size of the effect of gender on risk aversion was also numerically greater than that with the model-agnostic overall proportion bet measure, though this difference was not statistically significant (r = 0.14, transformed from d = 0.3 vs r = 0.18, transformed from d = 0.37: Steiger’s Z = 0.77, p = 0.22065). These results were unaffected by a sensitivity analysis in which participants whose data were not fit significantly better by our winning model than by chance were removed (Table S4).

### Model Parameters are Not Associated with Mental Health Symptoms

To assess whether model parameters were related to mental health symptoms, we calculated Pearson’s correlations between each pair of variables. However, we found no significant relationship between any of our model parameters and symptom scores, with all correlations r ≤ 0.07 (Table S3). Model-agnostic measures of task performance were also not related to mental health symptoms, with all correlations r ≤ 0.07 (Table S3). These results were unaffected by a sensitivity analysis in which participants whose data were not fit significantly better by our winning model than by chance were removed (Table S4). The distributions of questionnaire scores and model-agnostic measures are given in Table 2.

**Table 2.** Range of possible questionnaire scores or model-agnostic measures, along with mean, standard deviation, and median of participant scores. SRDS: Self-Rating Depression Scale, STAI-S: State Trait Anxiety Inventory – State, STAI-T: State Trait Anxiety Inventory – Trait, BIS-11: Barratt Impulsivity Scale, TEPS: Temporal Experience of Pleasure Scale. QDM: Quality of Decision Making, OPB: Overall Proportion Bet, RA: Risk Adjustment, DA: Delay Aversion.

| | Measure | Range | Mean ± SD, Median |
| --- | --- | --- | --- |
| Questionnaire | SRDS | 20 - 80 | 38.95 ± 10.25, 39 |
| | STAI-S | 20 - 80 | 36.97 ± 12.77, 35 |
| | STAI-T | 20 - 80 | 42.75 ± 13.26, 42 |
| | TEPS | 18 - 108 | 80.28 ± 11.44, 81 |
| | BIS-11 | 30 - 120 | 57.99 ± 9.73, 58 |
| Model-Agnostic Measure | QDM | 0 – 1 | 0.96 ± 0.07, 1.00 |
| | OPB | 0 – 1 | 0.54 ± 0.15, 0.56 |
| | RA | -2.4 – 7.2 | 2.43 ± 1.27, 2.44 |
| | DA | -0.9 – 0.9 | 0.07 ± 0.19, 0.04 |

## Discussion

In this study, we used a computational analysis of the CANTAB CGT to precisely investigate the mechanistic relationship between task behaviour, symptoms of mental health disorders and demographic variables. We fit five different models, including two novel ones, and found that a novel model in which betting strategies are not influenced by the number of current points, and uses an *inverse* power function in the loss domain, captures the characteristics of a large online dataset very well. Somewhat surprisingly, we found no significant relationships between model parameters and symptoms of mental health problems, but we did find robust associations between the risk aversion parameter and age and gender, such that older people and women were more risk averse. It is noteworthy that these relationships with demographic variables were stronger than those with raw outcome measures such as overall proportion bet, highlighting the added precision and mechanistic insight gained from this modelling analysis.

Our best-fitting model suggests a mechanistic explanation for how participants approach risky decisions when provided with explicit information about the amounts and probabilities that may be won or lost. Our novel model (Inverse Gains and Losses – IGL) varies from traditional risky decision-making models such as Prospect Theory in two main ways: 1) it assumes that participants’ betting strategies are independent of their current number of total points; and 2) it assumes that participants are consistently either risk averse or risk seeking across the domains of gains and losses. Both of these distinctions are likely to reflect the differences between tasks that Prospect Theory models were developed for and the CGT. The former were typically two-alternative forced choice tasks in which participants were required to choose between a risky gamble and a certain option (of winning or losing a number of points). As the current task asks participants to choose an amount to bet from five options, it involves a more fine-grained decision of how much one is willing to bet on a decision rather than just which option they choose. It is therefore likely to be more sensitive to characterising participants’ risk preferences. Furthermore, it is possible that because of the additional complexity in the gambling component (due to there being more options), the current number of points becomes a less salient component than the overarching bet choice strategy. Finally, previous literature suggests that on average participants tend to be risk averse in the gain domain but risk seeking in the loss domain (Kahneman & Tversky, 1979). Due to the power function in the gain domain, the inverse power function in the loss domain, and the values of the estimated risk aversion parameter in our winning model generally being below one, our model suggests that when performing the CGT participants tend to be risk averse in *both* domains. Again, this is likely to be due to key differences in the CGT compared to traditional risky decision-making tasks. Classical tasks are typically ‘additive’ in nature, such that the amount to be gained or lost is irrespective of the number of points the participant has. This contrasts with the CGT, where participants have to choose a bet from different proportions of their current number of points, which means that when following previous gains, participants have the potential to gain even more, often referred to as a ‘multiplicative’ feature. Multiplicative tasks can also often lead to bigger losses than additive tasks, and recent theories in ergodicity economics suggest that a multiplicative environment should therefore foster higher levels of risk aversion (Meder et al., 2021), which our results support.

Analysis of relationships between model parameters and symptoms of mental health problems revealed no significant findings. This result was somewhat surprising given the previous literature reporting more conservative risk attitudes in CGT in various groups with depressive symptoms (Mannie et al., 2015; Murphy et al., 2001; Rawal et al., 2013), as well as the previous algorithmic analysis of CGT in patients with substance use disorders (Romeu et al., 2020). However, our results are consistent with a recent study assessing CGT performance in an adolescent cohort (using standard model-agnostic measures) in which there was also no convincing evidence of a relationship between risk taking and depressive symptoms (Lewis et al., 2021), while clear relationships with age and gender were observed. It is possible that these inconsistencies are due to general population samples not capturing many participants at the more severe end of the symptom scales, and therefore limiting the size of any relationship. Alternatively, previous findings may have been due to chance or confounding variables. It is also possible that these models and parameters are not sufficiently sensitive to capture more specific differences in reward seeking or risk-taking behaviour seen in depression. For example, Tavares et al found that unmedicated depressed patients had worse quality of decision making in the colour choice component of the task specifically on trials that followed a loss (Taylor Tavares et al., 2007), which could be further explored with learning models. This highlights an interesting possible follow-up computational study for analysis of this task and its relationship with depression.

We found relationships between many model parameters and demographic variables of which two survived correction for multiple comparisons: older people and women were found to have higher levels of risk aversion. This finding has been consistently reported in previous literature with large cohort samples and model-agnostic CGT outcome measures such as overall proportion bet (Deakin et al., 2004; Lewis et al., 2021). Here, we replicated these findings with the behavioural measures, but further showed that the relationships between these demographic variables and risk aversion, a parameter from a model-based analysis, were stronger. This demonstrates that a modelling analysis of task behaviour can lead to more precise measurements, and also that these parameters are more mechanistic in nature than traditional model-agnostic behavioural measures. Future research should implement this computational approach in existing or novel CGT datasets to better understand previous findings.

There are some caveats to our study that merit comment. Our data was collected online, which is important for accessing large samples for research, but there is less control over the participants that volunteer to take part in online studies. This might lead to biased samples and spurious correlations. However, as the associations with demographic variables have been reported before, including in large cohort studies, these are less likely to have led to the results reported here. Another caveat of online data collection specific to this task, is the difficulty of drawing a sharp distinction between true impulsivity and distraction. It is possible that some participants select the earliest presented bet every time to finish the task as quickly as possible, or that some let the timer run out and are not engaging with the task properly. We attempted to get around this by removing a small percentage of individuals who always chose the first or last bets, and those whose data were not well fit by our winning model. This did not affect the results. However, it remains difficult to distinguish inattentive behaviour from poor decision making and impulsivity in online data collection.

In conclusion, we have presented a modelling analysis of CANTAB CGT in a novel, large dataset. This work highlights the added precision that computational models can provide to explore relationships with both demographic and mental health symptom variables.

## Acknowledgements

The authors thank the MRC and Cambridge Cognition Ltd. who funded this work jointly through an MRC iCASE studentship grant MR/R015759/1. Cambridge Cognition employ FC and completed the data collection in this research. We also thank the Neuroscience and Mental Health Group, Institute of Cognitive Neuroscience, University College London, London, United Kingdom and Alexandra Pike, PhD, Department of Psychology, University of York, York, United Kingdom, for their suggestions and troubleshooting advice during the creation of this article

## Footnotes

1 Left formal education before age 16, 2: Left formal education at age 16, 3: Left formal education at age 17-18, 4: Undergraduate degree or equivalent, 5: Master’s degree or equivalent, 6: PhD or equivalent.

2 Note that the degrees of freedom vary slightly as some outcome measures are incalculable from certain data. For instance, delay aversion is incalculable if participants never chose the majority box colour in at least one condition. In our data, one participant did not obtain a risk adjustment score, and two participants did not obtain a delay aversion score.

## References

Ackerman, J. P., McBee-Strayer, S. M., Mendoza, K., Stevens, J., Sheftall, A. H., Campo, J. v., & Bridge, J. A. (2015). Risk-sensitive decision-making deficit in adolescent suicide attempters. Journal of Child and Adolescent Psychopharmacology, 25(2), 109–113. https://doi.org/10.1089/CAP.2014.0041

Baek, K., Kwon, J., Chae, J. H., Chung, Y. A., Kralik, J. D., Min, J. A., Huh, H., Choi, K. M., Jang, K. I., Lee, N. bin, Kim, S., Peterson, B. S., & Jeong, J. (2017). Heightened aversion to risk and loss in depressed patients with a suicide attempt history. Scientific Reports, 7(1). https://doi.org/10.1038/S41598-017-10541-5

Bechara, A., Damasio, A. R., Damasio, H., & Anderson, S. W. (1994). Insensitivity to future consequences following damage to human prefrontal cortex. Cognition, 50(1–3), 7–15. https://doi.org/10.1016/0010-0277(94)90018-3

Bernoulli, D. (1738). Specimen theoriae novae de mensura sortis. Commentarii Academiae Scientiarum Imperialis Petropolitanae, 5, 175–192.

Cáceda, R., Nemeroff, C. B., & Harvey, P. D. (2014). Toward an understanding of decision making in severe mental illness. The Journal of Neuropsychiatry and Clinical Neurosciences, 26(3), 196–213. https://doi.org/10.1176/APPI.NEUROPSYCH.12110268

Charpentier, C. J., Aylward, J., Roiser, J. P., & Robinson, O. J. (2017). Enhanced Risk Aversion, But Not Loss Aversion, in Unmedicated Pathological Anxiety. Biological Psychiatry, 81(12), 1014–1022. https://doi.org/10.1016/j.biopsych.2016.12.010

Deakin, J., Aitken, M., Robbins, T., & Sahakian, B. J. (2004). Risk taking during decision-making in normal volunteers changes with age. Journal of the International Neuropsychological Society, 10(4), 590–598. https://doi.org/10.1017/S1355617704104104

Deserno, L., Schlagenhauf, F., & Heinz, A. (2016). Striatal dopamine, reward, and decision making in schizophrenia. Dialogues in Clinical Neuroscience, 18(1), 77–89. https://doi.org/10.31887/dcns.2016.18.1/ldeserno

Gard, D. E., Gard, M. G., Kring, A. M., & John, O. P. (2006). Anticipatory and consummatory components of the experience of pleasure: A scale development study. Journal of Research in Personality, 40(6), 1086–1102. https://doi.org/10.1016/J.JRP.2005.11.001

Halahakoon, D. C., Kieslich, K., O’Driscoll, C., Nair, A., Lewis, G., & Roiser, J. P. (2020). Reward-Processing Behavior in Depressed Participants Relative to Healthy Volunteers: A Systematic Review and Meta-analysis. JAMA Psychiatry, 77(12), 1286–1295. https://doi.org/10.1001/JAMAPSYCHIATRY.2020.2139

Hutton, S. B., Murphy, F. C., Joyce, E. M., Rogers, R. D., Cuthbert, I., Barnes, T. R. E., McKenna, P. J., Sahakian, B. J., & Robbins, T. W. (2002). Decision making deficits in patients with first-episode and chronic schizophrenia. Schizophrenia Research, 55(3), 249–257. https://doi.org/10.1016/S0920-9964(01)00216-X

Huys, Q. J. M., Cools, R., Gölzer, M., Friedel, E., Heinz, A., Dolan, R. J., & Dayan, P. (2011). Disentangling the roles of approach, activation and valence in instrumental and pavlovian responding. PLoS Computational Biology, 7(4), 1002028. https://doi.org/10.1371/journal.pcbi.1002028

Kahneman, D., & Tversky, A. (1979). Prospect Theory: An Analysis of Decision under Risk. Econometrica, 47(2), 263–291.

Kyte, Z. A., Goodyer, I. M., & Sahakian, B. J. (2005). Selected executive skills in adolescents with recent first episode major depression. Journal of Child Psychology and Psychiatry and Allied Disciplines, 46(9), 995–1005. https://doi.org/10.1111/J.1469-7610.2004.00400.X

Lewis, G., Srinivasan, R., Roiser, J., Blakemore, S. J., Flouri, E., & Lewis, G. (2021). Risk-taking to obtain reward: Sex differences and associations with emotional and depressive symptoms in a nationally representative cohort of UK adolescents. Psychological Medicine. https://doi.org/10.1017/S0033291720005000

Mannie, Z. N., Williams, C., Browning, M., & Cowen, P. J. (2015). Decision making in young people at familial risk of depression. Psychological Medicine, 45(2), 375–380. https://doi.org/10.1017/S0033291714001482

Meder, D., Rabe, F., Morville, T., Madsen, K. H., Koudahl, M. T., Dolan, R. J., Siebner, H. R., & Hulme, O. J. (2021). Ergodicity-breaking reveals time optimal decision making in humans. PLOS Computational Biology, 17(9), e1009217. https://doi.org/10.1371/JOURNAL.PCBI.1009217

Murphy, F. C., Rubinsztein, J. S., Michael, A., Rogers, R. D., Robbins, T. W., Paykel, E. S., & Sahakian, B. J. (2001). Decision-making cognition in mania and depression. Psychological Medicine, 693. https://doi.org/10.1017/S0033291701003804

Patton, J. H., Stanford, M. S., & Barratt, E. S. (n.d.). Factor Structure of the Barratt Impulsiveness Scale. https://doi.org/10.1002/1097-4679

Pedersen, A., Göder, R., Tomczyk, S., & Ohrmann, P. (2017). Risky decision-making under risk in schizophrenia: A deliberate choice? Journal of Behavior Therapy and Experimental Psychiatry, 56, 57–64. https://doi.org/10.1016/J.JBTEP.2016.08.004

Peterson, J. C., Bourgin, D. D., Agrawal, M., Reichman, D., & Griffiths, T. L. (2021). Using large-scale experiments and machine learning to discover theories of human decision-making. Science, 372(6547), 1209–1214. https://doi.org/10.1126/SCIENCE.ABE2629/SUPPL_FILE/ABE2629-PETERSON-SM.PDF

Pratt, D. N., Barch, D. M., Carter, C. S., Gold, J. M., Ragland, J. D., Silverstein, S. M., & MacDonald, A. W. (2021). Reliability and Replicability of Implicit and Explicit Reinforcement Learning Paradigms in People With Psychotic Disorders. Schizophrenia Bulletin, 47(3), 731–739. https://doi.org/10.1093/SCHBUL/SBAA165

Rawal, A., Collishaw, S., Thapar, A., & Rice, F. (2013). “The risks of playing it safe”: a prospective longitudinal study of response to reward in the adolescent offspring of depressed parents. Psychological Medicine, 43(1), 27–38. https://doi.org/10.1017/S0033291712001158

Rogers, R. D., Everitt, B. J., Baldacchino, A., Blackshaw, A. J., Swainson, R., Wynne, K., Baker, N. B., Hunter, J., Carthy, T., Booker, E., London, M., Deakin, J. F. W., Sahakian, B. J., & Robbins, T. W. (1999). Dissociable Deficits in the Decision-Making Cognition of Chronic Amphetamine Abusers, Opiate Abusers, Patients with Focal Damage to Prefrontal Cortex, and Tryptophan-Depleted Normal Volunteers: Evidence for Monoaminergic Mechanisms. Neuropsychopharmacology 1999 20:4, 20(4), 322–339. https://doi.org/10.1016/s0893-133x(98)00091-8

Romeu, R. J., Haines, N., Ahn, W. Y., Busemeyer, J. R., & Vassileva, J. (2020a). A computational model of the Cambridge gambling task with applications to substance use disorders. Drug and Alcohol Dependence, 206, 107711. https://doi.org/10.1016/J.DRUGALCDEP.2019.107711

Romeu, R. J., Haines, N., Ahn, W. Y., Busemeyer, J. R., & Vassileva, J. (2020b). A computational model of the Cambridge gambling task with applications to substance use disorders. Drug and Alcohol Dependence, 206, 107711. https://doi.org/10.1016/J.DRUGALCDEP.2019.107711

Rubinsztein, J. S., Fletcher, P. C., Rogers, R. D., Ho, L. W., Aigbirhio, F. I., Paykel, E. S., Robbins, T. W., & Sahakian, B. J. (2001). Decision-making in mania: a PET study. Brain : A Journal of Neurology, 124(Pt 12), 2550–2563. https://doi.org/10.1093/BRAIN/124.12.2550

Rubinsztein, J. S., Michael, A., Underwood, B. R., Tempest, M., & Sahakain, B. J. (2006). Impaired cognition and decision-making in bipolar depression but no “affective bias” evident. Psychological Medicine, 36(5), 629–639. https://doi.org/10.1017/S0033291705006689

Sachdev, P. S., & Malhi, G. S. (2005). Obsessive-compulsive behaviour: a disorder of decision-making. The Australian and New Zealand Journal of Psychiatry, 39(9), 757–763. https://doi.org/10.1080/J.1440-1614.2005.01680.X

Smoski, M. J., Lynch, T. R., Rosenthal, M. Z., Cheavens, J. S., Chapman, A. L., & Krishnan, R. R. (2008). Decision-making and risk aversion among depressive adults. Journal of Behavior Therapy and Experimental Psychiatry, 39(4), 567–576. https://doi.org/10.1016/J.JBTEP.2008.01.004

Spielberger, C. D. (1983). State-trait anxiety inventory for adults. https://www.scirp.org/(S(351jmbntvnsjt1aadkposzje))/reference/ReferencesPapers.aspx?ReferenceID=1861523

Talwar, A., Huys, Q., Cormack, F., & Roiser, J. (2021). A Hierarchical Reinforcement Learning Model Explains Individual Differences in Attentional Set Shifting. BioRxiv, 2021.10.05.463165. https://doi.org/10.1101/2021.10.05.463165

Taylor Tavares, J. v., Clark, L., Cannon, D. M., Erickson, K., Drevets, W. C., & Sahakian, B. J. (2007). Distinct Profiles of Neurocognitive Function in Unmedicated Unipolar Depression and Bipolar II Depression. Biological Psychiatry, 62(8), 917–924. https://doi.org/10.1016/J.BIOPSYCH.2007.05.034

von Neumann, J., & Morgenstern, O. (1944). Theory of Games and Economic Behavior. Princeton, NJ: Princeton University Press.

Wilson, R. C., & Collins, A. G. E. (2019). Ten simple rules for the computational modeling of behavioral data. ELife, 8. https://doi.org/10.7554/ELIFE.49547

Woodrow, A., Sparks, S., Bobrovskaia, V., Paterson, C., Murphy, P., & Hutton, P. (2019). Decision-making ability in psychosis: a systematic review and meta-analysis of the magnitude, specificity and correlates of impaired performance on the Iowa and Cambridge Gambling Tasks. Psychological Medicine, 49(1), 32–48. https://doi.org/10.1017/S0033291718002660

Zung WWK. (1965). A Self-Rating Depression Scale. Archives of General Psychiatry, 12(1), 63–70. https://doi.org/10.1001/ARCHPSYC.1965.01720310065008

