## Supplementary Information for "Individual Variation in Risky Decisions Is Related to Age and Gender but not to Mental Health Symptoms"

**Additional Models**

We built two further models not included in the main text, but specified here for further information:

***S1. Separate Gains and Losses (SGL)***

This model builds on the result from model 5 (IGL), using power functions for gains and losses but with separate parameters. It again assumes that participants subjectively value the bet proportions, $b_{t}$, independently from their current wealth$.$

$$U^{S}\left( b_{t},r_{t} \right)=\left\{ \begin{aligned} {b_{t}}^{\rho}, &if r_{t}=1 \\ -{b_{t}}^{\rho*}, &if r_{t}= -1 \end{aligned} \right.$$

(S1)

The parameters in this model were $\theta=\{\alpha,c,\rho,\rho*,\beta,\gamma\}$ where $\alpha$ is the colour choice determinism, $c$ the colour choice bias, $\rho$ the risk aversion for gains, $\rho*$ the risk aversion for losses, $\beta$ the delay aversion and $\gamma$ the bet choice determinism.

***S2. Separate Gains and Losses + Loss Aversion (SGL + LA)***

Our final model is similar to model S1 (SGL) with the addition of a multiplicative loss aversion term. It again assumes that participants subjectively value the bet proportions independently from their current wealth: $x$ = $\pm B.$

$$U^{L}\left( b_{t},r_{t} \right)=\left\{ \begin{aligned} {b_{t}}^{\rho}, &if r_{t}=1 \\ -\partial{b_{t}}^{\rho*}, &if r_{t}=-1 \end{aligned} \right.$$

(S2)

The parameters in this model were $\theta=\{\alpha,c,\rho,\rho*,\partial,\beta,\gamma\}$ where $\alpha$ is the colour choice determinism, $c$ the colour choice bias, $\rho$ the risk aversion for gains, $\rho*$ the risk aversion for losses, $\partial$ the loss aversion, $\beta$ the delay aversion and $\gamma$ the bet choice determinism.

Supplementary Figures & Tables

**Table S1.** Parameter ranges, distributions, and recovery for supplementary models.

| **Model** | **Parameter** | **Range** | **Mean ± SD, Median** | **Recovery** |
| --- | --- | --- | --- | --- |
| S1. Separate Gains and Losses | Colour choice determinism | 0 - $\infty$ | 7.96 ± 4.18, 8.53 | 0.57 |
|  | Colour choice bias | 0 - 1 | 0.50 ± 0.06, 0.50 | 0.48 |
|  | Risk aversion gains | 0 - $\infty$ | 2.35 ± 6.46, 0.22 | -0.03 |
|  | Risk aversion losses | 0 - $\infty$ | 1.62 ± 1.24, 1.24 | 0.90 |
|  | Delay aversion | -$\infty$ - $\infty$ | 0.04 ± 0.07, 0.02 | 0.89 |
|  | Bet choice determinism | 0 - $\infty$ | 39.66 ± 33.63, 33.42 | 0.67 |
| S2. Separate Gains and Losses + Loss Aversion | Colour choice determinism | 0 - $\infty$ | 8.33 ± 5.04, 8.46 | 0.65 |
|  | Colour choice bias | 0 - 1 | 0.50 ± 0.07, 0.50 | 0.51 |
|  | Risk aversion gains | 0 - $\infty$ | 1.24 ± 3.79, 0.37 | 0.47 |
|  | Risk aversion losses | 0 - $\infty$ | 1.80 ± 1.20, 1.43 | 0.81 |
|  | Loss aversion | 0 - $\infty$ | 1.53 ± 1.44, 1.18 | 0.46 |
|  | Delay Aversion | -$\infty$ - $\infty$ | 0.04 ± 0.08, 0.03 | 0.83 |
|  | Bet choice determinism | 0 - $\infty$ | 34.13 ± 22.85, 30.29 | 0.59 |

**Table S2.** Relationships between parameters and symptoms. Pearson’s correlations and p-values for relationship between symptom questionnaire scores and untransformed parameters of the best-fitting model (IGL). The bracketed value by each questionnaire gives the degrees of freedom for the corresponding analyses. SRDS: Self-Rating Depression Scale, STAI-S: State Trait Anxiety Inventory – State, STAI-T: State Trait Anxiety Inventory – Trait, BIS-11: Barratt Impulsivity Scale, TEPS: Temporal Experience of Pleasure Scale.

|  | | Parameters | | | | | |
| --- | --- | --- | --- | --- | --- | --- | --- |
|  |  | Colour choice determinism | Colour choice bias | Risk aversion | Loss aversion | Delay aversion | Bet choice determinism |
| Questionnaire | SRDS (746) | -0.03, 0.35 | 0.06, 0.12 | 0.06, 0.11 | -0.00, 0.90 | 0.04, 0.33 | -0.04, 0.29 |
|  | TEPST (745) | -0.05, 0.19 | -0.03, 0.39 | 0.02, 0.53 | -0.06, 0.12 | -0.03, 0.49 | -0.05, 0.18 |
|  | STAI-S (742) | -0.03, 0.40 | 0.06, 0.11 | 0.04, 0.23 | -0.02, 0.68 | 0.03, 0.42 | 0.03, 0.47 |
|  | STAI-T (742) | 0.02, 0.50 | 0.06, 0.09 | 0.02, 0.58 | -0.01, 0.69 | 0.01, 0.88 | 0.02, 0.55 |
|  | BIS-11 (741) | -0.07, 0.04 | 0.02, 0.52 | 0.02, 0.54 | -0.07, 0.06 | 0.06, 0.12 | -0.05, 0.21 |

**Table S3.** Relationships between model-agnostic measures and symptoms. Pearson’s correlations and p-values for relationship between symptom questionnaire scores and model-agnostic measures of CGT task performance. Bracketed values give the degrees of freedom for the analysis. SRDS: Self-Rating Depression Scale, STAI-S: State Trait Anxiety Inventory – State, STAI-T: State Trait Anxiety Inventory – Trait, BIS-11: Barratt Impulsivity Scale, TEPS: Temporal Experience of Pleasure Scale.

|  | | Model-Agnostic Measure | | | |
| --- | --- | --- | --- | --- | --- |
|  |  | Quality of Decision Making | Overall Proportion Bet | Risk Adjustment | Delay Aversion |
| Questionnaire | SRDS | -0.04, 0.31 (746) | -0.00, 0.96 (746) | 0.01, 0.70 (745) | 0.04, 0.24 (744) |
|  | TEPST | -0.03, 0.42 (745) | 0.07, 0.07 (745) | -0.05, 0.14 (744) | -0.05, 0.19 (743) |
|  | STAI-S | -0.04, 0.32 (742) | 0.01, 0.85 (742) | 0.01, 0.70 (741) | 0.04, 0.26 (741) |
|  | STAI-T | 0.01, 0.88 (742) | 0.01, 0.78 (742) | 0.05, 0.15 (741) | 0.03, 0.43 (741) |
|  | BIS-11 | -0.05, 0.17 (741) | 0.03, 0.39 (741) | -0.05, 0.21 (740) | 0.06, 0.09 (740) |

**Table S4.** Relationships between key variables from the sensitivity analysis. SRDS: Self-Rating Depression Scale, STAI-S: State Trait Anxiety Inventory – State, STAI-T: State Trait Anxiety Inventory – Trait, BIS-11: Barratt Impulsivity Scale, TEPS: Temporal Experience of Pleasure Scale.

| **Variable 1** | **Variable 2** | **Effect Size** | **p value** |
| --- | --- | --- | --- |
| Risk Aversion Parameter | Age | r = -0.25 | 4.99x10^-12^ |
| Risk Aversion Parameter | Gender | Cohen’s d = 0.38 | 4.02x10^-7^ |
| Overall Proportion Bet | Age | r = -0.15 | 3.84x10^-5^ |
| Overall Proportion Bet | Gender | Cohen’s d = 0.32 | 2.29x10^-5^ |
| Risk Aversion Parameter | SDRS | r = 0.05 | 0.14 |
| Risk Aversion Parameter | TEPS | r = 0.02 | 0.59 |
| Loss Aversion Parameter | SDRS | r = 0.00 | 0.93 |
| Loss Aversion Parameter | TEPS | r = -0.05 | 0.14 |
| Overall Proportion Bet | SDRS | r = -0.01 | 0.88 |
| Overall Proportion Bet | TEPS | r = 0.07 | 0.07 |

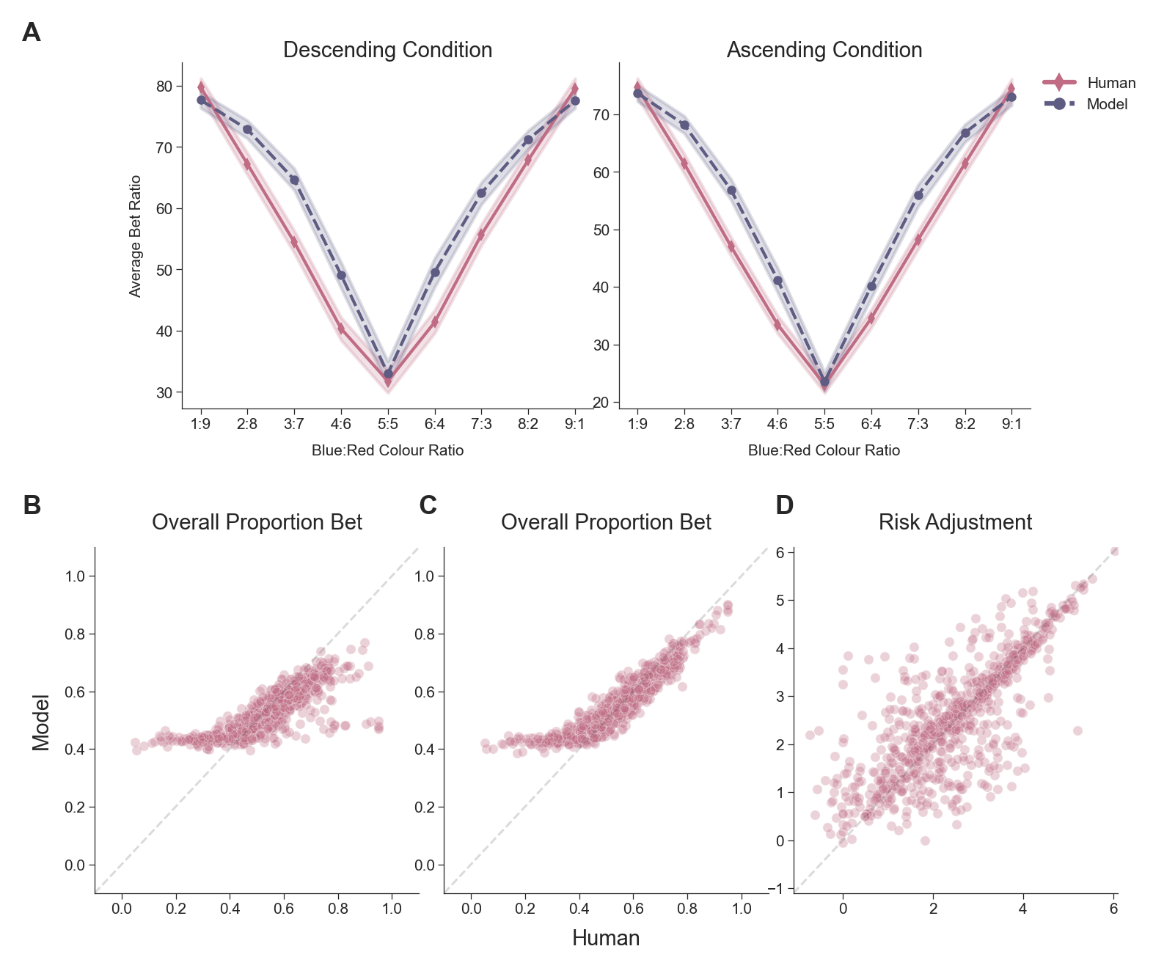

**Figure S1.** Model Weaknesses.

**A.** Average Group Fit of Prospect Theory Models. Human and model-simulated patterns of proportion bet averaged over subjects for each box ratio and task condition. Points indicate mean whilst bands represent 95% confidence intervals. **B.** Individual Differences in Overall Proportion Bet for Expected Utility Theory Model. Scatter plot of human vs model-simulated scores, r = 0.80. **C.** Individual Differences in Overall Proportion Bet for Expected Utility Fixed Gains Model. Scatter plot of human vs model-simulated scores, r = 0.92. **D.** Individual Differences in Risk Adjustment for Linear Loss Aversion Model. Scatter plot of human vs model-simulated scores, r = 0.77. The correlation between human and model data was calculated for 10 different simulations, and the average of these is reported here.
